# High throughput profiling of the B cell repertoire identifies systematic changes in the repertoire of individuals with Crohn’s disease

**DOI:** 10.1101/2025.09.01.673486

**Authors:** Aya K.H. Mahdy, Zahra Taheri, Marte Lie Høivik, Andre Franke, Hesham ElAbd

## Abstract

The B cell repertoire contains the recombined DNA sequences that encodes the entire antibody repertoire of an individual. The repertoire is made from three antigenic binding chains, namely the immunoglobulin heavy chain (IGH) and two immunoglobulin light chains, κ (IGK) and λ (IGL). Compared to the T cell repertoire, the B cell repertoire is understudied in inflammatory bowel diseases (IBD) even though different antibodies such as ASCA and ANCA have been shown to be elevated in individuals with IBD. Furthermore, most IBD B cell repertoire studies have profiled the repertoire of treated individuals, thus capturing the combined effect of treatment and disease on the repertoire. To address this limitation, we profiled the repertoire of 24 treatment-naive individuals with CD with matching 24 symptomatic controls. The repertoire of individuals with CD showed a significant reduction in diversity and an increase in clonality, suggesting an antigen-driven expansion of clonotypes that might be driving the disease. Furthermore, we observed a significant reduction in the expansion of IgM and IgD and an expansion of IgA2 and IgG2 clonotypes in individuals with CD relative to controls. Lastly, we observed a reduction in the somatic hypermutation rate in the IGH J gene, particularly in IgM and IgA1 clonotypes, among individuals with CD relative to controls. Thus, despite the small sample size, we identified multiple alterations in the B cell repertoire of individuals with CD, highlighting the potential of the B cell repertoire in identifying antigenic exposures implicated in the diseases, demanding now larger international studies ideally including also treatment-naive and pre-clinical cases.

## Introduction

Inflammatory bowel disease (IBD) is an incurable immune-mediated inflammatory disease that predominantly affects the gastrointestinal tract (GIT). It is observed clinically in two main forms: Crohn’s disease (CD), which is characterized by patchy transmural inflammation of different sections of the GIT, predominantly the ileum and the colon, and ulcerative colitis (UC), which is restricted to the colon. The exact cause(s) of IBD remain unknown, however, different genetic variants have been associated with the disease, such as *ATG16L*^*1,2*^, and *NOD2*^*3*^. Additionally, several human leukocyte antigen (HLA) alleles have been implicated in IBD, such as HLA-DRB1*03:01^*4*^, or in a specific subset of individuals with ileal CD, such as HLA-DRB1*07:01^5^ and HLA-DRB1*15:01 in UC^6^. Besides genetic predispositions, other risk factors have been identified, *e.g*., smoking, microbial dysbiosis^7^, antibiotic intake^8^, and previous episodes of infectious mononucleosis^9^, which is mainly caused by an Epstein-Barr virus (EBV) infection.

From an immunological perspective, different alterations and dysregulated processes have been identified in individuals with IBD, including dysregulated responses toward the gut microbiome^10^ and mycobiome^11^. We previously observed a significant expansion of a subset of type II natural killer T cells, termed Crohn’s-associated invariant T (CAIT) cells, in individuals with CD^12,13^. Elevated antibody responses against bacterial flagellins^14^ and several human herpesviruses, predominantly EBV have been reported by others^15^. Recently, we also performed a large-scale analysis of the T cell repertoire of individuals with IBD in comparison to matching controls, identifying thousands of clonotypes that were significantly expanded in individuals with IBD^16,17^. These disease-associated clonotypes represent an immunological fingerprint for common antigenic exposures implicated in the disease. Nonetheless, these antigens remain to be identified as T cell repertoire sequencing enables the identification of clonotypes, *i.e*., V(D)J recombination sequences forming the T cell receptor, and not the exact antigens presented by HLA proteins and recognized by these T cells^18^.

The dependency of T cells on HLA proteins, which are highly polymorphic^19^, renders analysing the T cell repertoire a challenging task, specifically in case-control studies where thousands of samples are needed to pinpoint specific T cells involved in the disease across different HLA context^16,17,20^. Thus, identifying B cells recognizing the same antigens across multiple individuals with CD might require a smaller sample size as B cell receptors (BCRs) recognise their antigenic targets directly independent of any presenting molecules, *e.g*. HLA proteins. Relative to the T cell repertoire, which has been studied by others^21–24^ and us^12,16,20^, the B cell repertoire of individuals with IBD remains under investigated.

The B cell repertoire is a composite of all immunoglobulins heavy and light chains present in a sample, such as peripheral blood. These chains are generated from somatic recombination processes termed V(D)J recombination that generate chains with enormous sequence diversity. This diversity is also augmented by a unique process that happens in B cells, termed somatic hypermutation (SHM), where random single-nucleotide polymorphisms (SNPs) are introduced into the formed immunoglobulin chains to increase their affinity toward a particular antigen. The Ig heavy chain (IGH) has five main isotypes, namely, μ, δ, ε, α, and γ, which form the IgM, IgD, IgE, IgA, and IgG, respectively. Furthermore, the α and γ isotypes have different subclasses, namely, α1 and α2 that encode IgA1 and IgA2, and γ1, γ2, γ3, and γ4, which encode the IgG1, IgG2, IgG3, and IgG4 subtypes. There are two light chains, κ (IGK) and λ (IGL), but they only have one constant region and do not undergo class-switching.

Besides the seminal study by Bashford-Rogers *et al*. ^25^ and colleagues, which compared the BCR repertoire of six different autoimmune diseases, not many studies have investigated the immune repertoire in individuals with CD or UC. Scheid *et al*.^26^ profiled the B cell repertoire of colonic tissues from individuals with UC and showed a bias from IgA1 and IgA2 isotype usage toward IgG2 usage in inflamed tissues. Similarly, Chen and colleagues^27^ profiled the B cell repertoire of multiple tissues in individuals with IBD and showed dysregulated B cell responses in individuals with CD. Lastly, Kotagiri *et al*.^*28*^ also profiled the repertoire of individuals with IBD and healthy controls and identified BCRs shared among individuals with CD.

While these studies have advanced our understanding of B cell responses driving the disease and put us one-step closer toward identifying the antigen underlying the disease, they have been conducted on treated individuals and thus they capture the effect of the disease as well as medications on the repertoire. Here, we aimed at investigating the B cell receptor of 24 treatment-naive individuals with CD and matching symptomatic controls from the Norwegian inception cohort IBSEN-III^29^.

## Results

### The B cell repertoire of treatment-naive individuals with CD exhibits systematic differences in clonality and diversity

We started our analysis by comparing the clonal expansion landscape of individuals with CD relative to SCs. The expansion-rank distributions of the IGH, IGK, and IGL repertoires of individuals with CD differed from that of matching SCs (**Fig. 1A-1C**). This indicates systematic differences in the repertoire, specifically regarding the expansion and the evenness of the repertoire. To follow this further, we compared the Shannon-diversity between individuals with CD and SCs (**Methods**) which, across the three antigenic binding chains, showed a significant reduction in diversity (**Fig. 1D**). The same finding was also observed when comparing the Simpson diversity which is less sensitive to rarer clonotypes, among the repertoire of the three antigenic binding chains (**Fig. 1E**). Lastly, we compared the evenness of the repertoire by comparing the Gini-inequality index (**Methods)** in individuals with CD relative to SC which showed a significant increase in the inequality, *i.e*. a decrease in evenness, in the repertoire of individuals with CD across the three chains (**Fig. 1F**). These findings indicate an antigen driven expansion in the B cell repertoire of individuals with CD relative to symptomatic controls.

**Figure 1:**
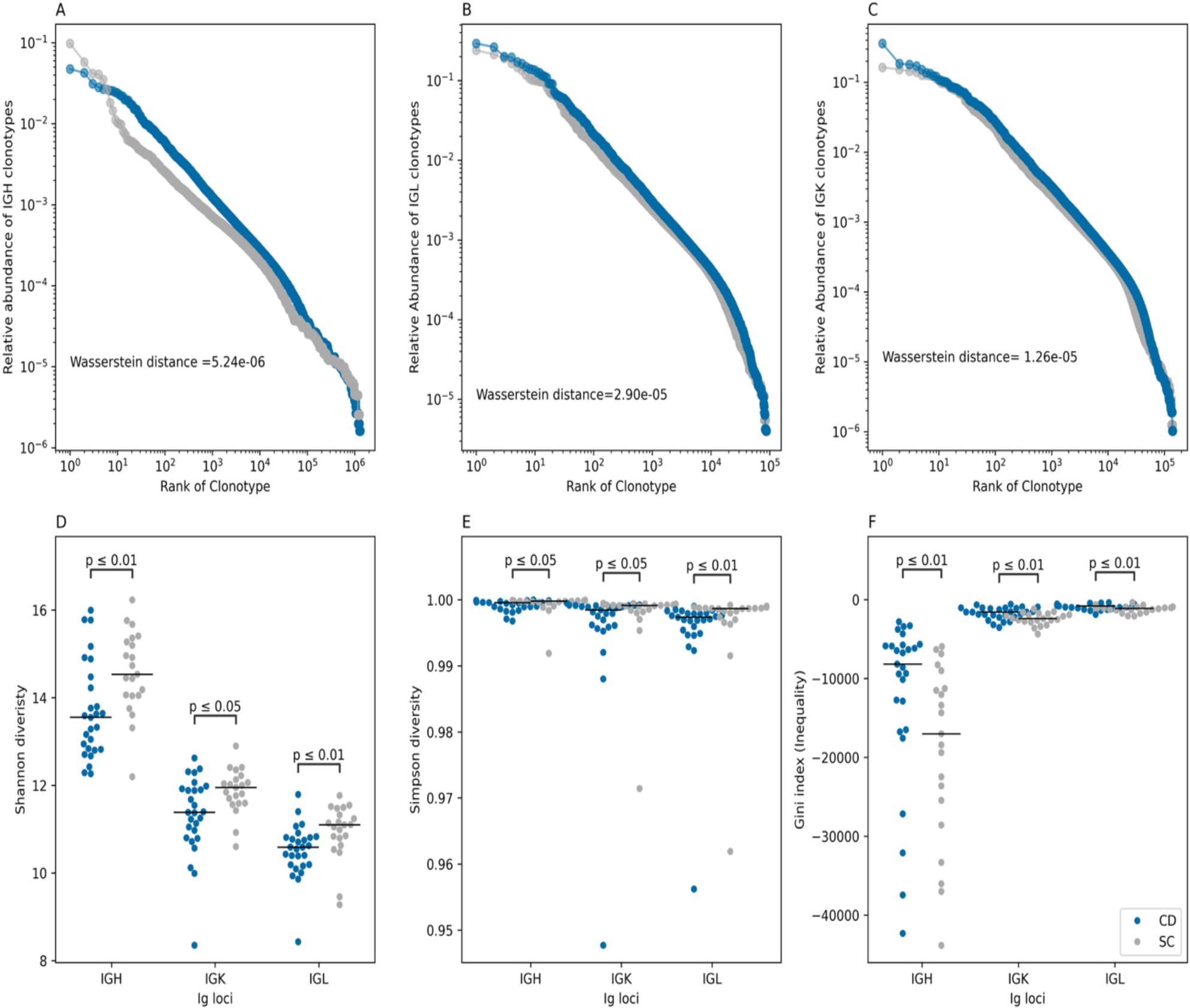
The BCR repertoire of individuals with CD is significantly different from that of matched symptomatic controls. (**A**-**C**) shows the abundance-rank distribution of the IGH (**A**), IGK (**B**), and IGL (**C**) repertoire of individuals with CD and SC. In panels (**A**-**C**), the distribution of individuals with CD was compared to that of controls using Wasserstein distance between the two distributions. (**D**) depicts the reduction in the Shannon diversity of the BCR repertoire in individuals with CD relative to SCs, while (**E**) depicts the reduction in Simpson diversity in individuals with CD. Lastly, (**F**) depicts the increase in inequalities in the repertoire of individuals with CD relative to controls. In panels (**D**-**E**), the different properties of the repertoire were compared using a two-sided Mann-Whitney U test.

### The BCR repertoire encodes personalized antigenic exposure trajectories

Motivated by these findings, we aimed at identifying expanded B cell clonotypes that might be driving the disease. Thus, we extracted the most expanded 250 clonotypes across the entire cohort and investigated their occurrence patterns among individuals with CD and SC. The IGH repertoire was exceptionally diverse relative to the IGK and IGL repertoire, where expanded clones were only detected in an individual and were not shared across individuals (**Fig. S1**). Expanded IGK and IGL clonotypes were more public and were detected among different individuals, however, there was no clear clustering based on disease status (**Fig. S2**-**S3**). Subsequently, we expanded our analysis to the entire repertoire, not only expanded clonotypes and calculating the Jaccard index between each unique pair of samples. Across the three chains, limited overlap was observed among the different study participants, furthermore, there was no clear clustering pattern based on disease status (**Fig. S4**-**S6**).

### CD does not alter the VDJ recombination landscape of the blood B cell receptor repertoire

Next, we aimed at quantifying the impact of CD on the abundance of different V and J gene combinations. Within the IGH repertoire, some VJ recombination pairs dominated the repertoire, for example, *IGHV3-23*/*IGHJ4* derived clonotypes (**Fig. S7**). Similar findings were detected at the IGK repertoire, where *IGK3-20* paired with *IGKJ1* dominated the repertoire (**Fig. S8**), and the IGL repertoire, where *IGL2-14* paired with *IGLJ2* formed a high percentage of the IGL repertoire (**Fig. S9**) in individuals with and without CD. These findings indicate that CD induces a change in repertoire diversity and clonality, but it is not large enough to change the landscape of V(D)J recombination, at least not at a magnitude detected with our study’s statistical power. To extend this analysis to other less frequent VJ combinations, we compared the expansion of each unique VJ gene combination between individuals with and without CD (**Methods**). Nonetheless, after correcting for multiple testing, we could not detect any significant VJ gene combination that was significantly expanded in CD relative to SC across any of the three loci (**Fig. S10**).

### Individuals with CD have a significant reduction in the expansion of IgM and IgD clonotypes and an expansion of IgA2 and IgG2 clonotypes

Subsequently, we aimed at investigating isotype chain usage by individuals with and without CD (**Methods**). We observed a significantly lower levels of IgM and IgD in individuals with CD relative to SC (**Fig. 2A**), similarly, we observed a significant increase in the levels of IgA2 and IgG2 in individuals with CD relative to SC (**Fig. 2A**). This effect mostly persisted after removing the effect of expansion, *i.e*., counting each clonotype as a singleton (**Fig. 2B**). Thus, it shows that in CD there is a decrease in the number and expansion levels of IgM and an increase in the number and expansion levels of IgA2 and IgG2 clonotypes. We also observed a significant reduction in the expansion levels of IgD in individuals with CD. To study this further, we aimed at investigating the class-switching flux between the different IGH isotypes (**Methods**). There was an overall comparable level of flux between CD and SC, where most class-switching events were observed between IgM and IgA1, followed by IgM to IgA2, then IgM and IgG1, and lastly, IgM and IgG2 (**Fig. S11-S12**).

**Figure 2:**
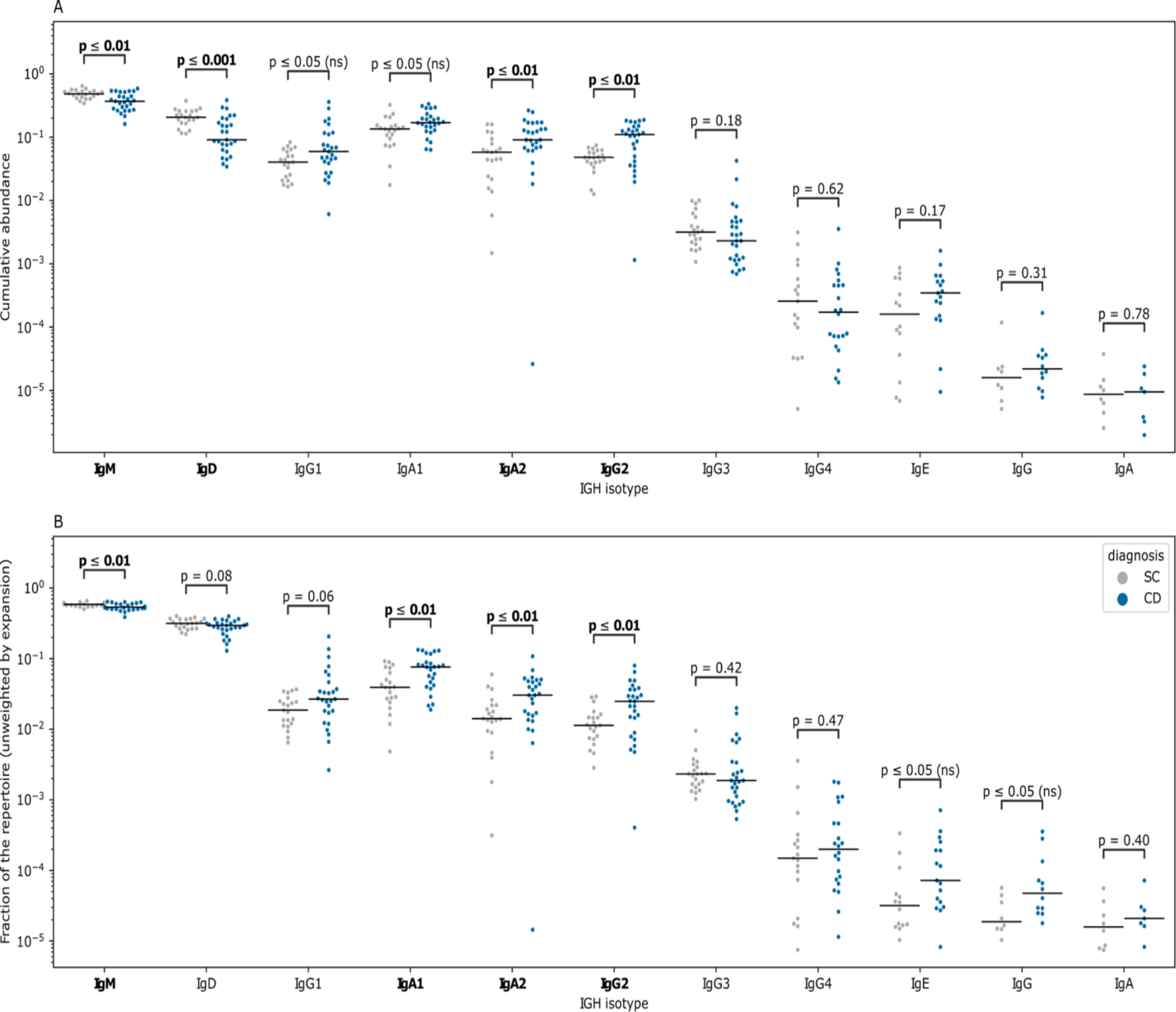
Alterations in the IGH isotype utilization in individuals with CD relative to controls. (**A**) The cumulative abundance is weighted by the expansion of each clonotype, among the different isotypes and subclasses of the IGH chains. (**B**) depicts the same relationship but without considering the expansion of each clonotype; that is, the count of each clonotype over the total number of counts defined in each sample was used to compute the fractions depicted in (**B**). In both panels, isotype utilization was compared using a two-sided Mann-Whitney U test, and the results were corrected for multiple testing using the Benjamin-Hochberg false discovery rate approach. Statistically significant results are highlighted in bold type.

### Individuals with CD have a reduction in the rate of somatic hypermutation in the IGHJ gene but no the IGHV gene

Finally, we aimed to quantify the effect of CD on the overall rates of SHM (**Methods**). We started by investigating the rate of SHM in the IGHV and IGHJ genes among the different loci independent of the disease, where we observed an overall higher level of SHM rates on all IgA and IgG isotypes relative to IgM (**Fig. S13**). This is consistent with our immunological understanding of class-switching and affinity maturation, where initial responses are generated via IgM, which then class switch to other types such as IgA or IgG. Hence, it is expected that the rate of SHMs will be higher in class-switched antibodies relative to IgM. After that, we compared the rate of SHMs between individuals with CD and SC, where repeated antigenic exposures with the same antigen might result in elevated levels of SHMs in individuals with CD. Across the different IGHV genes and isotypes, there was an overall comparable level of SHMs (**Fig. 3A**), nonetheless, there was a significant increase in the rate of SHM in the IGHJ gene (**Fig. 3B**).

**Figure 3:**
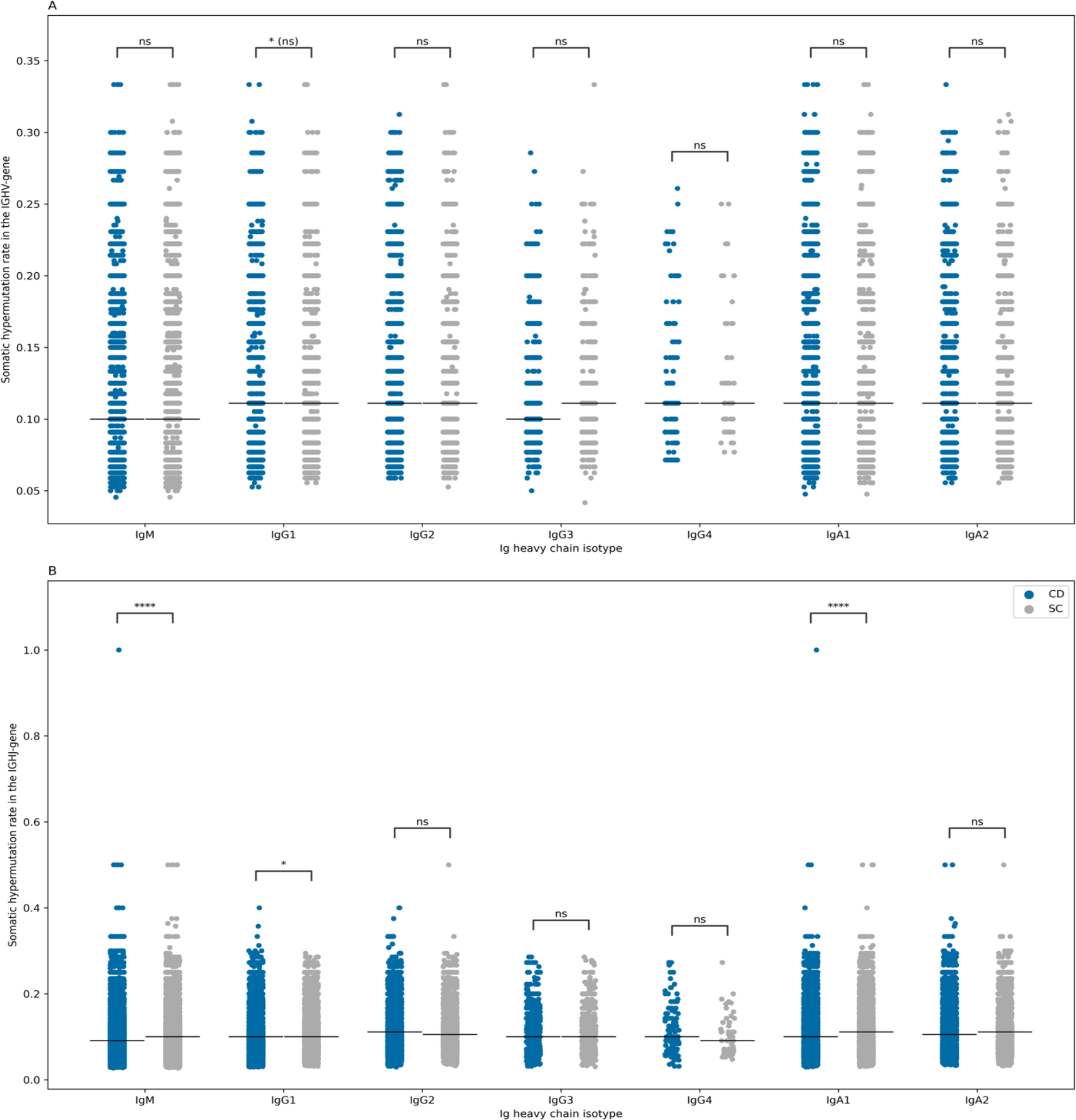
The difference in the rate of somatic hypermutation (SHM) in the V gene (**A**) and J gene (**B**) among the different isotypes of the IGH loci and between individuals with CD and SC. In both panels, isotype utilization was compared using a two-sided Mann-Whitney U test, and the results were corrected for multiple testing using the Benjamin-Hochberg false discovery rate approach with *(ns) indicate nominal significance that was not significant after multiple testing correction.

## Discussion

Our investigation of the BCR repertoire in treatment-naive individuals has validated previous findings discovered in treated individuals. These include the significant reduction in the expansion of IgM and IgD in the repertoire of individuals with CD relative to SC, reported by Bashford-Rogers *et al*.^*25*^. In addition, we also validated the higher expansion of IgA2 in individuals with CD and reported multiple alterations in the IgG isotype utilisation for example and an increase in the expansion of IgG2 clonotypes in individuals with CD. IgM and IgD are primarily utilized by naive B cells and effector B cells prior to class switching. Hence, this might indicate a reduction in the amount of naive B cells present in the repertoire of individuals with CD relative to SC as a result of higher antigenic exposure, either because of a persistent infection and/or a leakier gut barrier. This is also supported by the higher levels of IgA, particularly IgA2, and IgG2, that are more expanded in the blood repertoire of individuals with CD relative to SC^25^. We also observed a significant reduction in the SHM rate in the J gene of individuals with CD relative to SC, particularly in IgM and IgA, suggesting a less-strict class switching process.

In conclusion, our study identified multiple alterations in the repertoire of treatment-naive individuals with CD, highlighting treatment-independent and disease-specific effects on the B cell repertoire of individuals with CD. Future studies shall aim at increasing the sample size of included individuals and focusing the analysis on antigen-exposed memory B cells to identify antigenic exposures driving the disease.

## Methods

### Cohort description

The cohort contains treatment-naive individuals with CD or symptomatic controls, *i.e*., individuals with the symptoms of IBD but without endoscopic or radiological findings supporting their diagnosis. The study participants were recruited as a part of the Inflammatory Bowel Disease in South-Eastern Norway III (IBSEN III) study^29^. The IBSEN-III study was approved by the South-Eastern Regional Committee for Medical and Health Research Ethics (Ref 2015/946-3) and performed in accordance with the Declaration of Helsinki.

### Profiling the B cell receptor

RNA was extracted from PAXgene tubes collected from study participants using a QIAcube (Qiagen). After quality controls, 300ng of RNA were used to profile the IGH and IGK repertoire of the samples using the Human IG RNA Multiplex kits from MiLaboratories^®^ according to the manufacturer’s instructions. After library preparation, Illumina’s unique dual indexing adaptors were used to tag each cell with a unique index, then samples were pooled together, cleaned with magnetic beads (AMPure XP, Beckman Coulter). Next, samples were sequenced using a 250bp X2 using a NextSeq 1000 at the Illumina Solutions Center in Berlin. After sequencing and samples demultiplexing, the generated reads of each sample were processed using MiXCR^30^ to identify clonotypes present of each sample. After clonotype identification, non-productive clonotypes were filtered out as we focused the analysis on functional, *i.e*., productive immunoglobulin chains.

### Diversity and inequality calculations

The Shannon diversity of each repertoire was calculated according to (**Eq.1**), where R represents the repertoire size, *i.e*. the number of clonotypes in a repertoire, while the *expansion* _*i*_ represents the clonal expansion of the i^th^ clonotype in the repertoire.

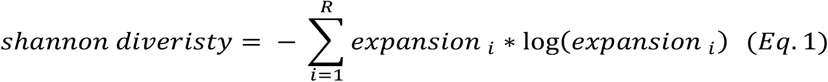

Similarly, Simpson diversity index was calculated according to (**Eq. 2**), which is similar to Shannon diversity, except that there is no weighting by the logarithm of expansion, *i.e*. log(*expansion* _*i*_).

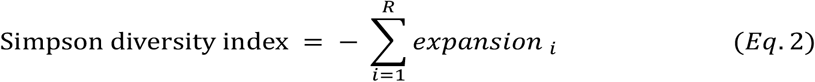

Lastly, the Gini inequality coefficient was calculated according to (**Eq. 3**)

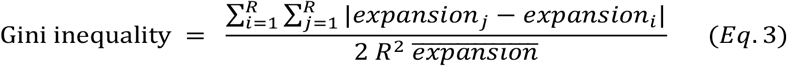

Where R is the repertoire size, 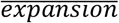 representing the mean value of expression across all clonotypes. Finally, the *expansion*_*i*_ and *expansion*_*i*_ represent the expansion of the i^th^ and j^th^ clonotype with the repertoire of clonotypes identified in the input sample.

### Comparing the VJ recombination frequency among the different conditions

To compare the frequency of different VJ recombination between individuals with CD and SC, we calculated the mean of each combination in all samples using (**Eq. 4**)

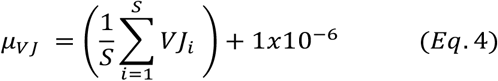

Where μ_*VJ*_represents the mean of expression for a specific VJ recombination, S represents the number of samples in a specific group, *e.g*., CD. The 1*x*10^−6^ represent a pseudo-count for a VJ recombination to accommodate for samples where a particular VJ was not detected. After that, the fold change between the group was calculated using (**Eq. 5**), where the mean expression of each VJ was compared between CD and SCs. Lastly, the P-value was calculated using a two-sided Mann–Whitney U test.

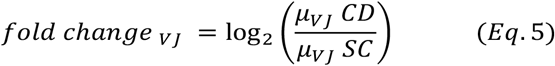

### Isotype flux

To calculate the flux between different isotypes, we extract clonotypes that had the same V-J and CDR3 amino acid but with different isotypes, for example, IgM and IgA. Given that the same V(D)J rearrangement has been observed at both isotypes, then we assume that this is due to a class-switching event, for example, IgM to IgG or an IgM to IgA. Lastly, we divided by the total number of transitions in a sample to get the normalized flux of a specific isotype.

To represent this mathematically, we introduced the set V, which represents the set of allowed class switching, Here. we considered the following transitions: *V* = {(*I gM*→*I gA1)*, (*I gM*→*I gA2)*,(*I gM*→*I gG*1),(*I gM*→*I gG*2),(*I gM*→*I gG*3),(*I gM*→*I gG4)*}. Across each sample *s* and condition, *c* we calculated the transition for a specific pair in V as 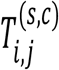 which represents the total number of transitions between an allowed set of isotypes from set V in a sample *s* and a condition *c*. Lastly, we divide the number of transitions between the i^th^ and j^th^ isotypes by the total number of transitions across all isotypes to come up with the normalized flux between isotypes in a given sample as, depicted in (**Eq. 6**).

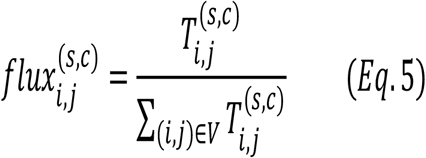

### Somatic hypermutation analysis

To calculate the rate of somatic hypermutations (SHMs), we first identified somatic hypermutations in the V and J genes by comparing their sequence in given clonotypes to their germline sequences. After that we divided the number of identified mutations by the length of germline genes to calculate the rate of SHM in a specific clonotype. Here, we only focused on protein-coding, that is, non-synonymous mutations.

## Ethical approval

The IBSEN III study was approved by the South-Eastern Regional Committee for Medical and Health Research Ethics (Ref 2015/946-3) and performed in accordance with the Declaration of Helsinki. All patients signed an informed consent form prior inclusion in this study, and the data were stored in services for sensitive data (TSD) at the University of Oslo.

## Availability of Data

Data are not deposited in a public repository due to data privacy regulations in Norway and our institution. However, data are available upon request, if the aims of the planned analyses are covered by the written informed consent signed by the participants, pending an amendment to the ethical approvals and a material & data transfer agreement between the institutions.

## References

1. Hampe J., Franke A., Rosenstiel P., Till A., Teuber M., Huse K., et al. A genome-wide association scan of nonsynonymous SNPs identifies a susceptibility variant for Crohn disease in ATG16L1. Nat Genet 2007;39(2):207–11. Doi: 10.1038/ng1954.

2. Lavoie S., Conway KL., Lassen KG., Jijon HB., Pan H., Chun E., et al. The Crohn’s disease polymorphism, ATG16L1 T300A, alters the gut microbiota and enhances the local Th1/Th17 response. Elife 2019;8:e39982. Doi: 10.7554/eLife.39982.

3. Hampe J., Cuthbert A., Croucher PJP., Mirza MM., Mascheretti S., Fisher S., et al. Association between insertion mutation in <em>NOD2</em> gene and Crohn’s disease in German and British populations. The Lancet 2001;357(9272):1925–8. Doi: 10.1016/S0140-6736(00)05063-7.

4. Goyette P., Boucher G., Mallon D., Ellinghaus E., Jostins L., Huang H., et al. High-density mapping of the MHC identifies a shared role for HLA-DRB1*01:03 in inflammatory bowel diseases and heterozygous advantage in ulcerative colitis. Nat Genet 2015;47(2):172–9. Doi: 10.1038/ng.3176.

5. Goyette P., Boucher G., Mallon D., Ellinghaus E., Jostins L., Huang H., et al. High-density mapping of the MHC identifies a shared role for HLA-DRB1*01:03 in inflammatory bowel diseases and heterozygous advantage in ulcerative colitis. Nat Genet 2015;47(2):172–9. Doi: 10.1038/ng.3176.

6. Degenhardt F., Mayr G., Wendorf M., Boucher G., Ellinghaus E., Ellinghaus D., et al. Trans-ethnic analysis of the human leukocyte antigen region for ulcerative colitis reveals shared but also ethnicity-specific disease associations. Hum Mol Genet 2021. Doi: 10.1093/hmg/ddab017.

7. Shan Y., Lee M., Chang EB. The gut microbiome and inflammatory bowel diseases. Annu Rev Med 2022;73(1):455–68.

8. Faye AS., Allin KH., Iversen AT., Agrawal M., Faith J., Colombel J-F., et al. Antibiotic use as a risk factor for inflammatory bowel disease across the ages: a population-based cohort study. Gut 2023;72(4):663. Doi: 10.1136/gutjnl-2022-327845.

9. Ebert AC., Harper S., Vestergaard M V., Mitchell W., Jess T., Elmahdi R. Risk of inflammatory bowel disease following hospitalisation with infectious mononucleosis: nationwide cohort study from Denmark. Nat Commun 2024;15(1):8383. Doi: 10.1038/s41467-024-52195-8.

10. Uchida AM., Boden EK., James EA., Shows DM., Konecny AJ., Lord JD. Escherichia coli–Specific CD4+ T Cells Have Public T-Cell Receptors and Low Interleukin 10 Production in Crohn’s Disease. Cell Mol Gastroenterol Hepatol 2020;10(3):507–26. Doi: 10.1016/j.jcmgh.2020.04.013.

11. Martini GR., Tikhonova E., Rosati E., DeCelie MB., Sievers LK., Tran F., et al. Selection of cross-reactive T cells by commensal and food-derived yeasts drives cytotoxic TH1 cell responses in Crohn’s disease. Nat Med 2023;29(10):2602–14. Doi: 10.1038/s41591-023-02556-5.

12. Rosati E., Martini GR., Pogorelyy M V., Minervina AA., Degenhardt F., Wendorf M., et al. A novel unconventional T cell population enriched in Crohn’s disease. Gut 2022;71(11):2194 LP – 2204. Doi: 10.1136/gutjnl-2021-325373.

13. Mahdy A., ElAbd H., Kriukova V., Olbjørn C., Perminow G., Bengtson MB., et al. P0125 Crohn’s-associated invariant T Cells are associated with disease severity and location and are not afected by medication intake. J Crohns Colitis 2025;19(Supplement_1):i507–8. Doi: 10.1093/ecco-jcc/jjae190.0299.

14. Bourgonje AR., Andreu-Sánchez S., Vogl T., Hu S., Vich Vila A., Gacesa R., et al. Phage-display immunoprecipitation sequencing of the antibody epitope repertoire in inflammatory bowel disease reveals distinct antibody signatures. Immunity 2023;56(6):1393-1409.e6. Doi: 10.1016/j.immuni.2023.04.017.

15. Nandy A., Petralia F., Porter C., Elledge S., Anand R., Croitoru K., et al. Epstein-Barr Virus (EBV) Exposure Precedes Crohn’s Disease Development. Gastroenterology 2025. Doi: 10.1053/j.gastro.2025.01.247.

16. Pesesky M., Bharanikumar R., Le Bourhis L., ElAbd H., Rosati E., Carty CL., et al. Antigen-driven expansion of public clonal T cell populations in inflammatory bowel diseases. J Crohns Colitis 2025:jjaf048. Doi: 10.1093/ecco-jcc/jjaf048.

17. Elabd H., Mahdy A., Franke A. OP11 Analysing the T cell receptor beta chain repertoire of 2,800 Inflammatory Bowel Disease patients identifies public T cell responses involved in the pathogenesis of Crohn’s Disease and Ulcerative Colitis and quantifies the impact of surgery and therapy on the immune repertoire of IBD patients. J Crohns Colitis 2025;19(Supplement_1):i21–3. Doi: 10.1093/ecco-jcc/jjae190.0011.

18. Mahdy AKH., Lokes E., Schöpfel V., Kriukova V., Britanova O V., Steiert TA., et al. Bulk T cell repertoire sequencing (TCR-Seq) is a powerful technology for understanding inflammation-mediated diseases. J Autoimmun 2024;149:103337. Doi: 10.1016/j.jaut.2024.103337.

19. ElAbd H., Mahdy A., Wacker EM., Gretsova M., Ellinghaus D., Franke A. Decoding the restriction of T cell receptors to human leukocyte antigen alleles using statistical learning. BioRxiv 2025:2022–5.

20. ElAbd H., Pesesky M., Innocenti G., Chung BK., Mahdy AKH., Kriukova V., et al. T and B cell responses against Epstein–Barr virus in primary sclerosing cholangitis. Nat Med 2025. Doi: 10.1038/s41591-025-03692-w.

21. Allez M., Auzolle C., Ngollo M., Bottois H., Chardiny V., Corraliza AM., et al. T cell clonal expansions in ileal Crohn’s disease are associated with smoking behaviour and postoperative recurrence. Gut 2019;68(11):1961–70.

22. Pesesky M., Carty CL., Singh N., Le Bourhis L., Rosati E., Bokemeyer B., et al. DOP47 Identification and characterization of T-cell receptor sequences associated with Crohn’s Disease. J Crohns Colitis 2022;16(Supplement_1):i096–7. Doi: 10.1093/ecco-jcc/jjab232.086.

23. Chapman CG., Yamaguchi R., Tamura K., Weidner J., Imoto S., Kwon J., et al. Characterization of T-cell Receptor Repertoire in Inflamed Tissues of Patients with Crohn’s Disease Through Deep Sequencing. Inflamm Bowel Dis 2016;22(6):1275–85. Doi: 10.1097/MIB.0000000000000752.

24. Doorenspleet ME., Westera L., Peters CP., Hakvoort TBM., Esveldt RE., Vogels E., et al. Profoundly Expanded T-cell Clones in the Inflamed and Uninflamed Intestine of Patients With Crohn’s Disease. J Crohns Colitis 2017;11(7):831–9. Doi: 10.1093/ecco-jcc/jjx012.

25. Bashford-Rogers RJM., Bergamaschi L., McKinney EF., Pombal DC., Mescia F., Lee JC., et al. Analysis of the B cell receptor repertoire in six immune-mediated diseases. Nature 2019;574(7776):122–6. Doi: 10.1038/s41586-019-1595-3.

26. Scheid JF., Eraslan B., Hudak A., Brown EM., Sergio D., Delorey TM., et al. Remodeling of colon plasma cell repertoire within ulcerative colitis patients. Journal of Experimental Medicine 2023;220(4):e20220538. Doi: 10.1084/jem.20220538.

27. Chen D., Xu S., Li S., Wang Q., Li H., He D., et al. The multi-organ landscape of B cells highlights dysregulated memory B cell responses in Crohn’s disease. Natl Sci Rev 2025;12(4):nwaf009. Doi: 10.1093/nsr/nwaf009.

28. Kotagiri P., Rae WM., Bergamaschi L., Pombal D., Lee J-Y., Noor NM., et al. Disease-specific B cell clones are shared between patients with Crohn’s disease. Nat Commun 2025;16(1):3689. Doi: 10.1038/s41467-025-58977-y.

29. Kristensen VA., Opheim R., Perminow G., Huppertz-Hauss G., Detlie TE., Lund C., et al. Inflammatory bowel disease in South-Eastern Norway III (IBSEN III): a new population-based inception cohort study from South-Eastern Norway. Scand J Gastroenterol 2021;56(8):899–905.

30. Bolotin DA., Poslavsky S., Mitrophanov I., Shugay M., Mamedov IZ., Putintseva E V., et al. MiXCR: software for comprehensive adaptive immunity profiling. Nat Methods 2015;12(5):380–1. Doi: 10.1038/nmeth.3364.

