## Supplementary figures for "High throughput profiling of the B cell repertoire identifies systematic changes in the repertoire of individuals with Crohn’s disease"

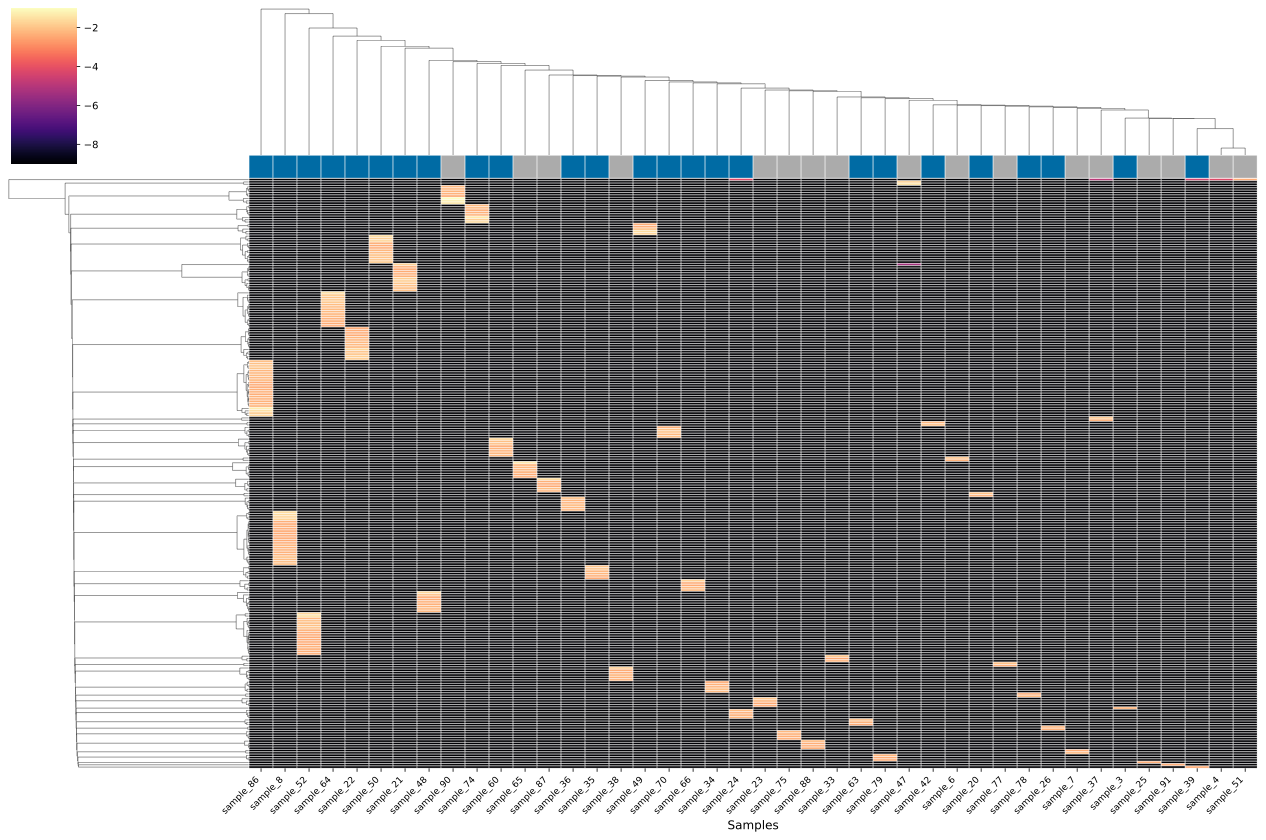

**Figure S1**: the expansion of the top 250 most expanded IGH clonotypes in individuals with CD (depicted in blue) and SC (depicted in grey). Most of these clonotypes appear to be sample-specific and are not shared among any of the study participants. The colour bar shows the log transformation of the abundance of each clonotype in a specific sample (shown on the x-axis).

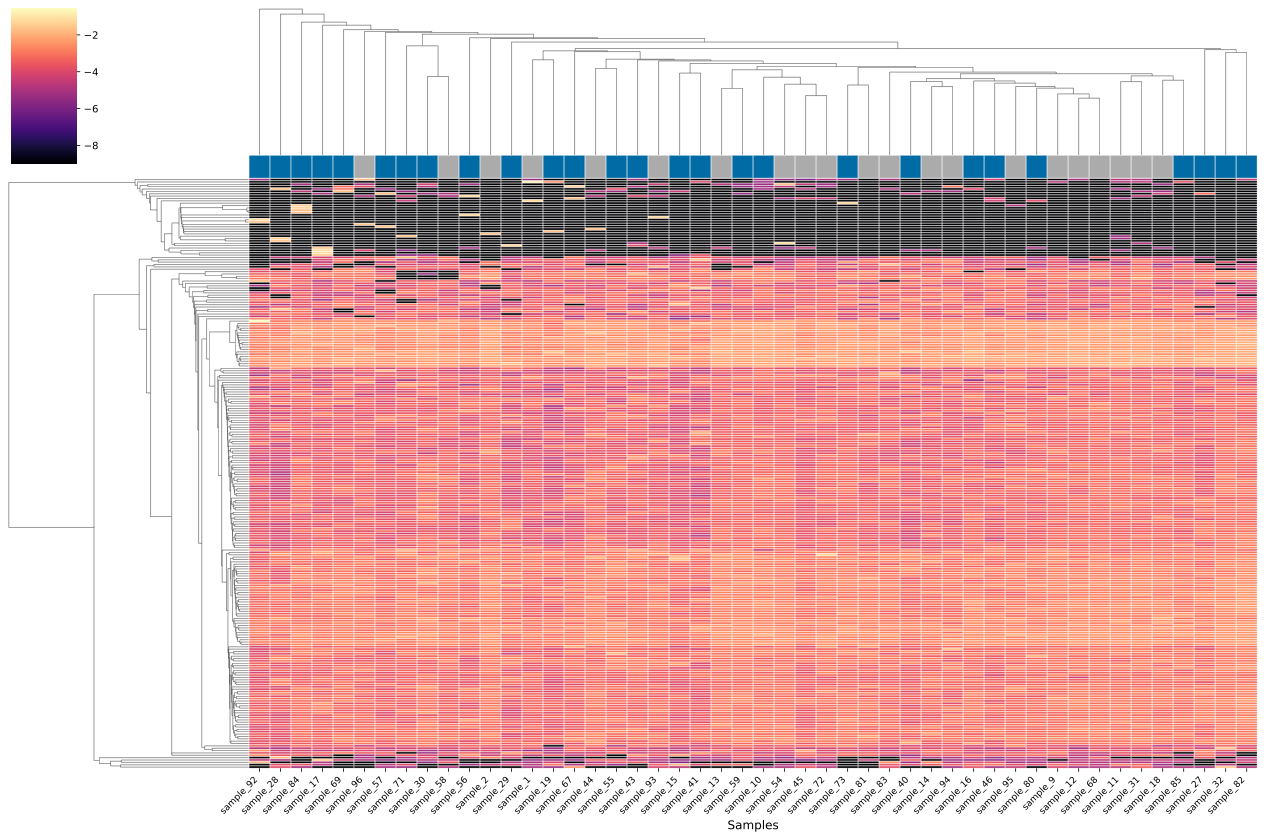

**Figure S2**: The expansion of the top 250 most expanded IGK clonotypes in individuals with CD (depicted in blue) and SC (depicted in grey). Most of these clonotypes appear to be highly public and are shared among most, if not all, study participants; however, they lack a clear clustering pattern with the disease.

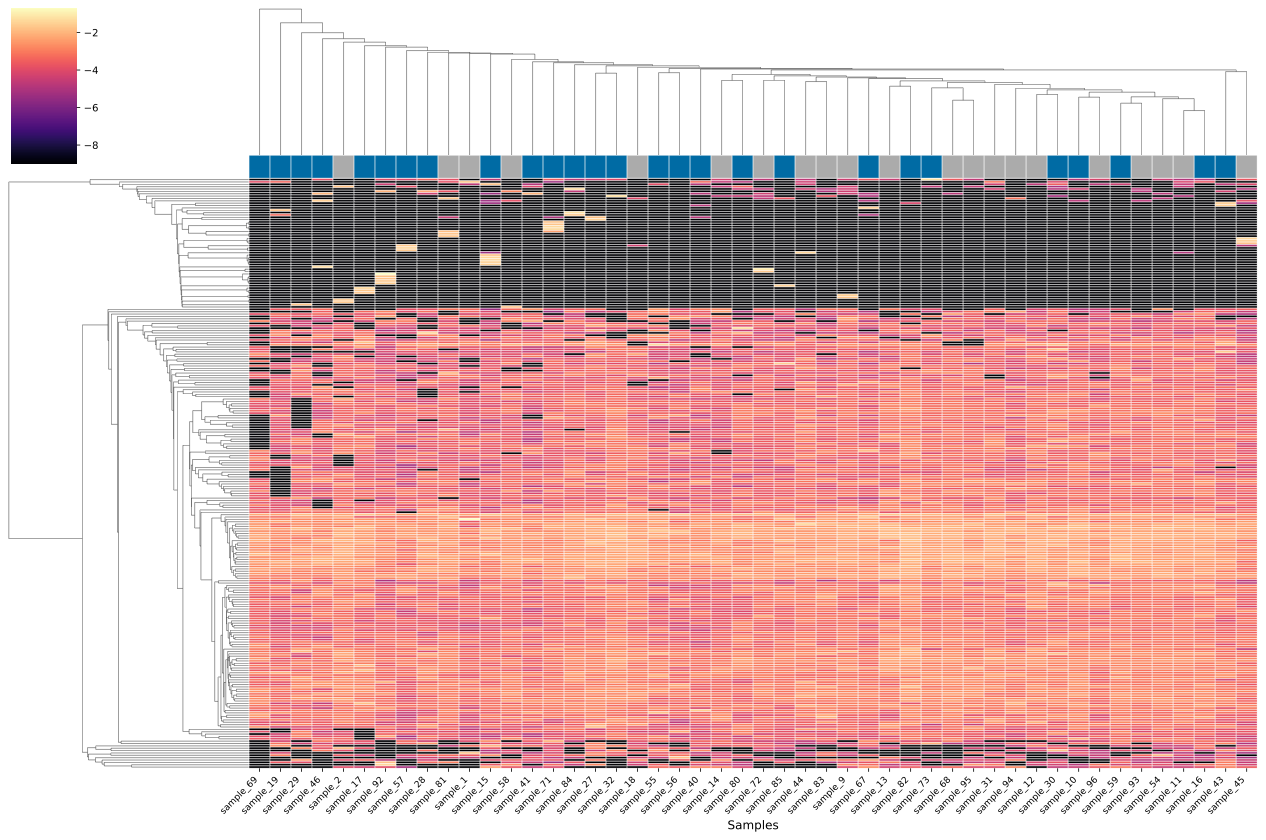

**Figure S3**: The expansion of the top 250 most expanded IGL clonotypes in individuals with CD (depicted in blue) and SC (depicted in grey). Most of these clonotypes appear to be highly public and are shared among most, if not all, study participants; however, they lack a clear clustering pattern with the disease.

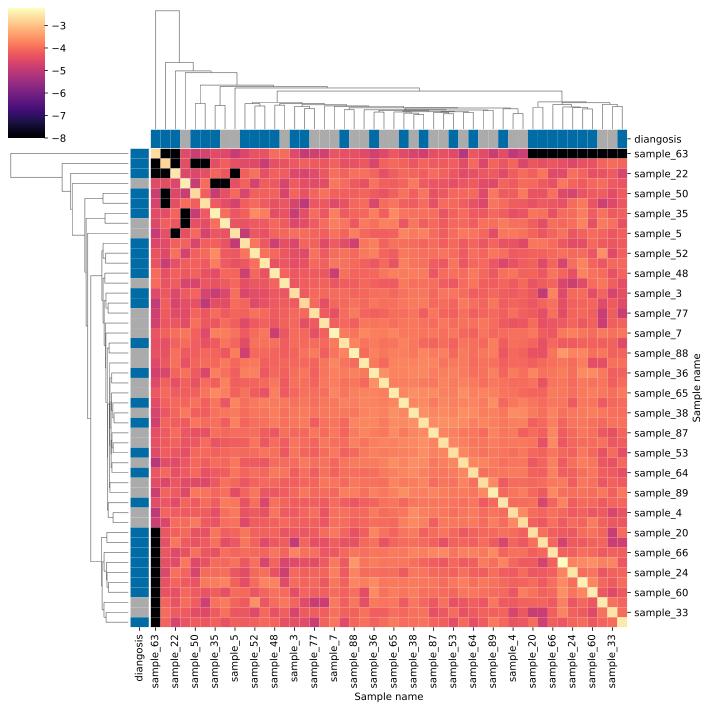

**Figure S4**: The overlap in the IGH repertoire of individuals with CD and SC as calculated using the Jaccard index. The color bar represents a log transformation of the Jaccard index between two samples.

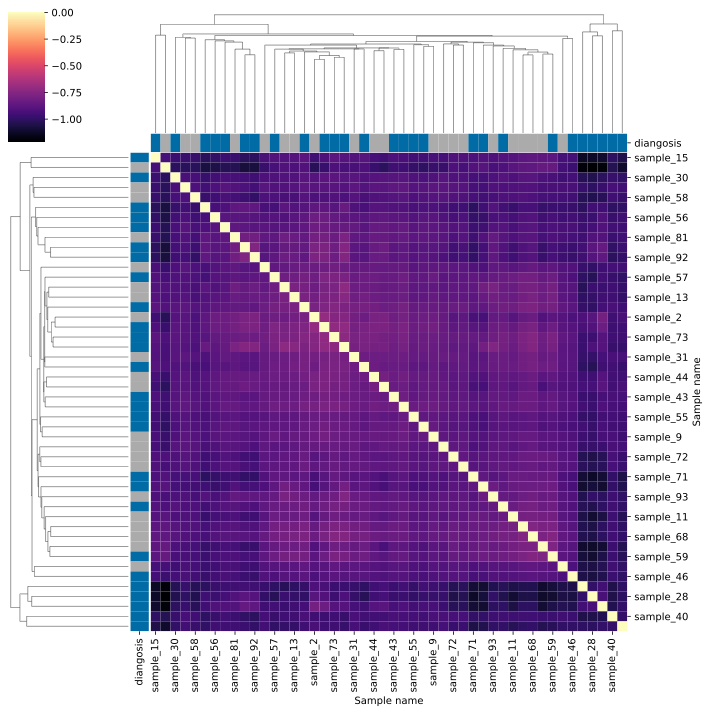

**Figure S5**: The overlap in the IGK repertoire of individuals with CD and SC as calculated using the Jaccard index. The color bar represents a log transformation of the Jaccard index between two samples.

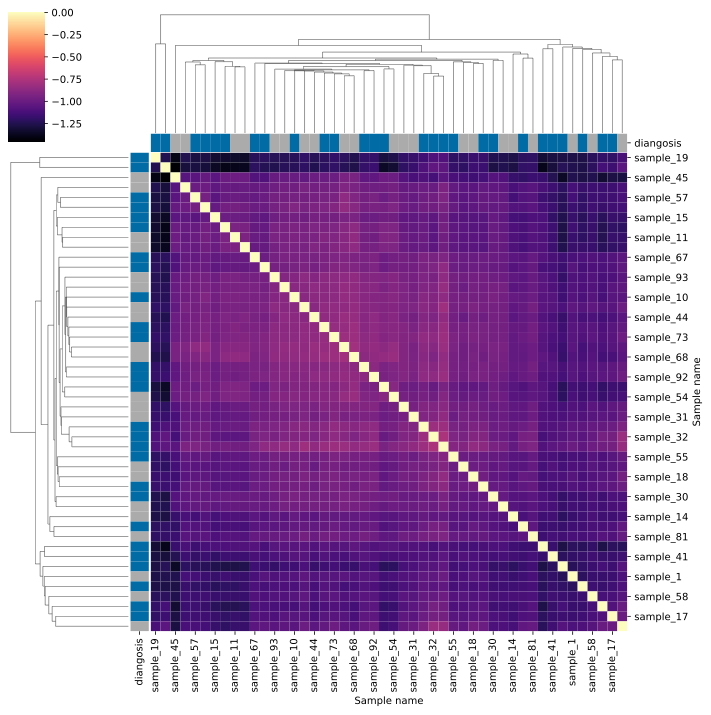

**Figure S6**: The overlap in the IGL repertoire of individuals with CD and SC as calculated using the Jaccard index. The color bar represents a log transformation of the Jaccard index between two samples.

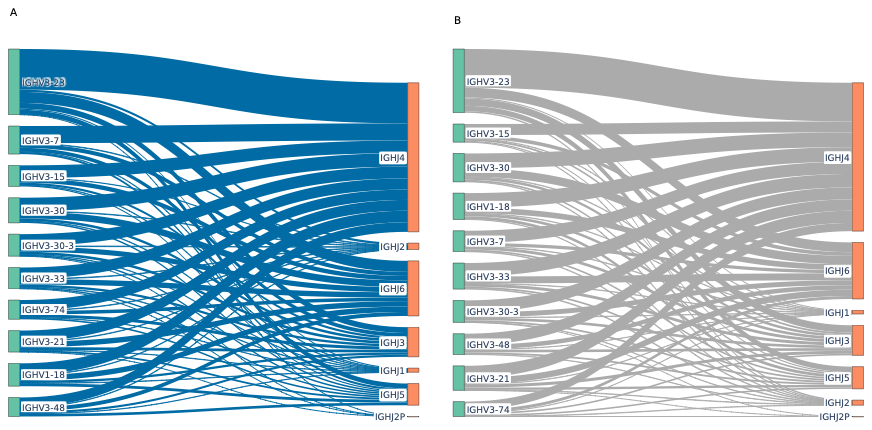

**Figure S7**: A representation of the most common VJ recombination observed across the IGH repertoire of individuals with CD (**A**) or SC (**B**).

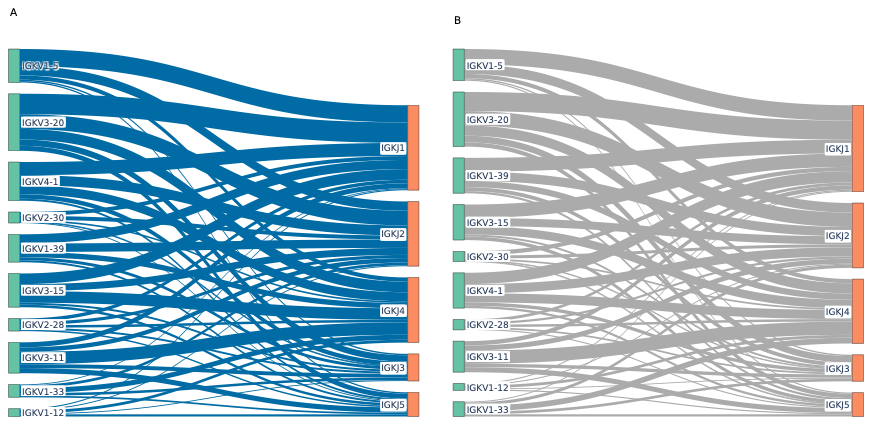

**Figure S8**: A representation of the most common VJ recombination observed across the IGK repertoire of individuals with CD (**A**) or SC (**B**).

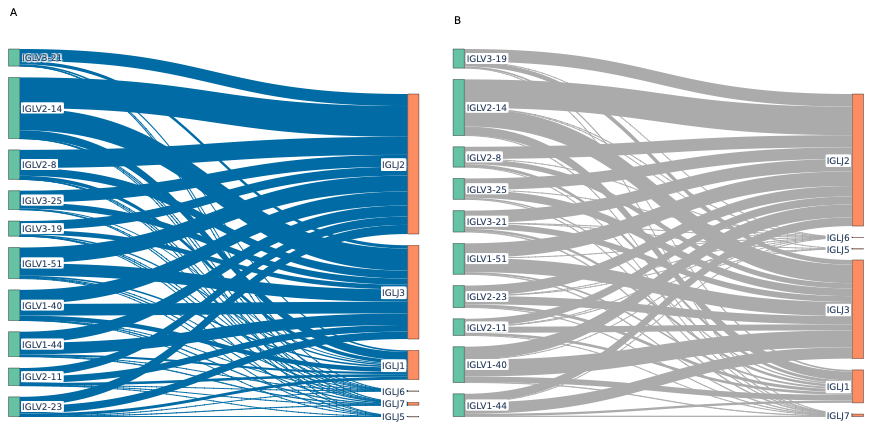

**Figure S9**: A representation of the most common VJ recombination observed across the IGL repertoire of individuals with CD (**A**) or SC (**B**).

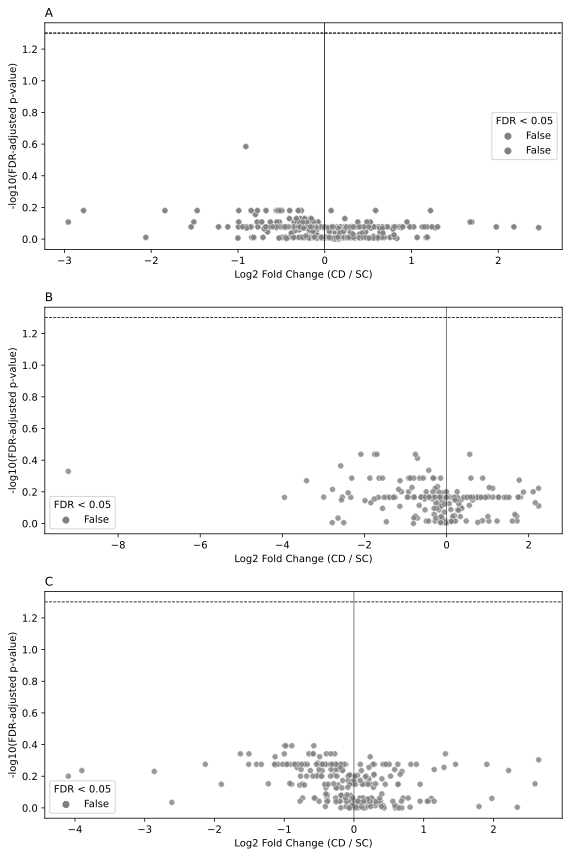

**Figure S10**: A volcano plot for the differential expansion of different VJ recombination in the IGH repertoire (**A**), IGL repertoire (**B**), or IGK repertoire (**C**). None of the tested recombination acrossany of the loci was significant after correcting for multiple testing.

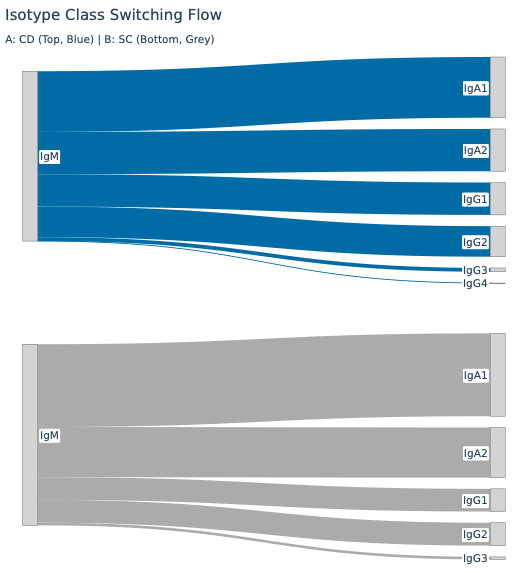

**Figure S11**: An isotype flux analysis showing the most common class-switches across the IGH repertoire of individuals with CD (**A**) and SC (**B**), with an overall similarity in which the most common translation was IgM to IgA1 or IgA2, followed by IgG1 and IgG2 in both phenotypes.

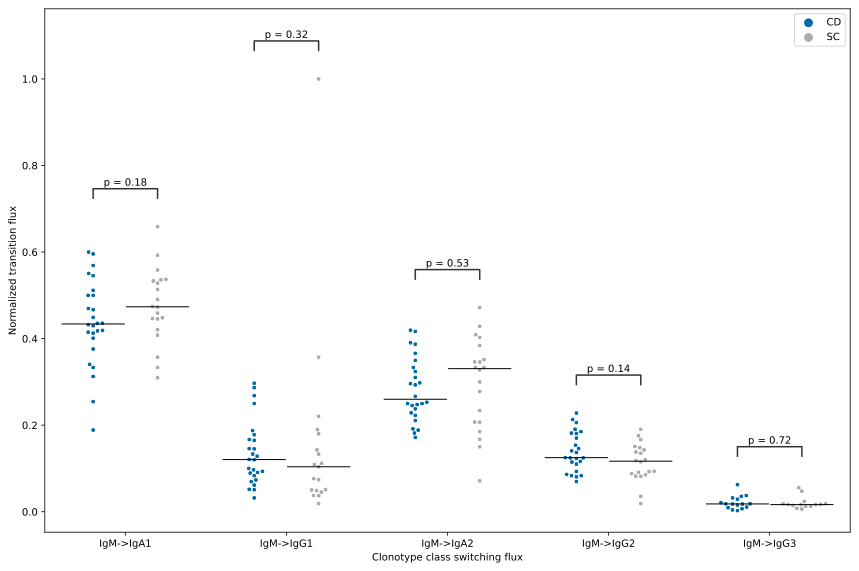

**Figure S12**: A comparison between the normalized flux of isotype switching between individuals with CD and SC, where no significant difference in the flux was observed between the two repertoires. The normalized flux was compared using a two-sided Mann-Whitney U test with correction for multiple testing.

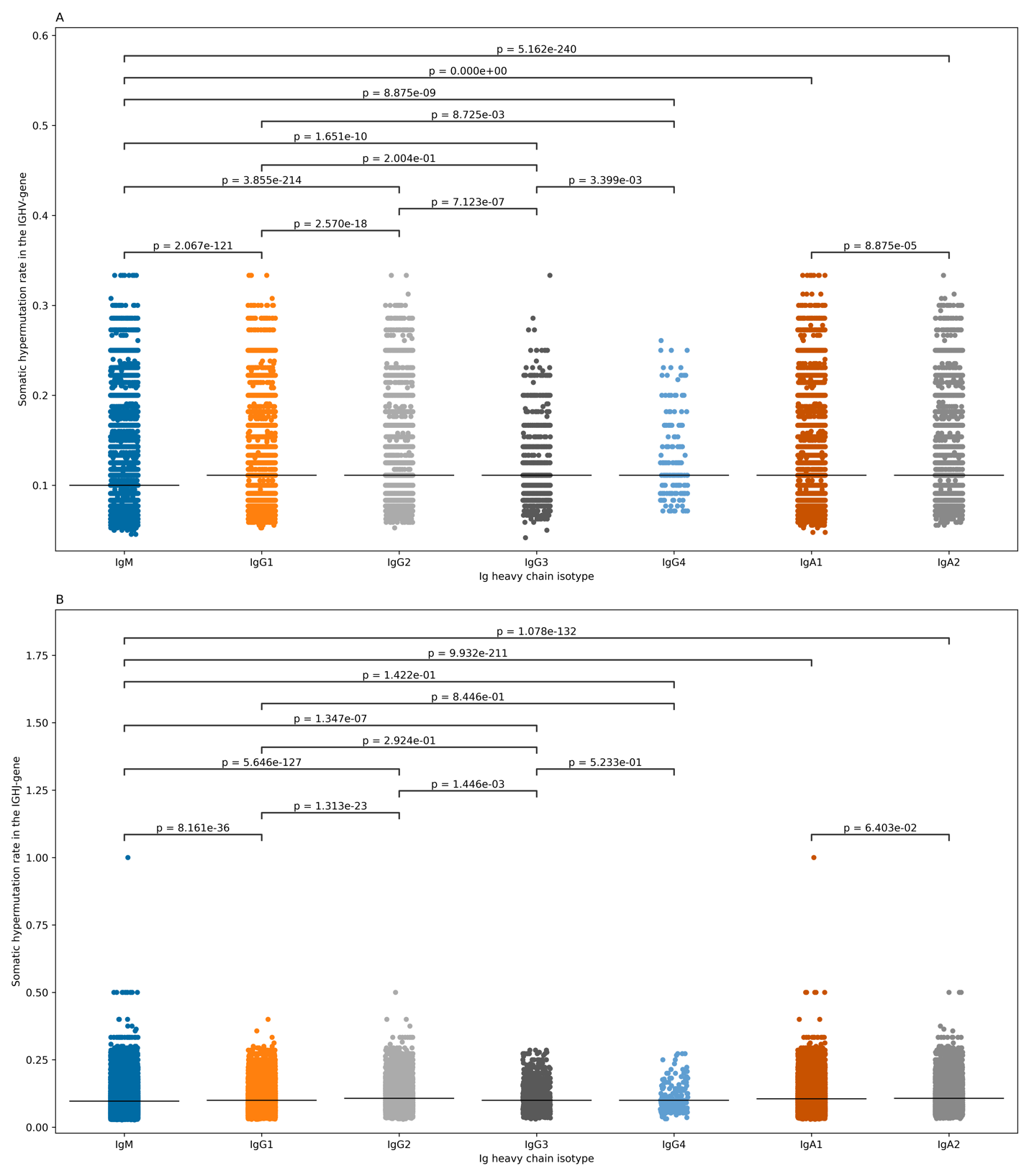

**Figure S13**: The difference in the rate of somatic hypermutation (SHM) in the V gene (**A**) and J gene (**B**) among the different isotypes of the IGH loci. The rate of SHMs were compared using a two-sided Mann-Whitney U test, and the results were corrected for multiple testing using the Benjamin-Hochberg false discovery rate approach.
